# Structural connectome topology relates to regional BOLD signal dynamics in the mouse brain

**DOI:** 10.1101/085514

**Authors:** Sarab S. Sethi, Valerio Zerbi, Nicole Wenderoth, Alex Fornito, Ben D. Fulcher

**Author notes:** These authors contributed equally to this work (Sarab S. Sethi, Valerio Zerbi).

## Abstract

Brain dynamics are thought to unfold on a network determined by the pattern of axonal connections linking pairs of neuronal populations; the so-called connectome. Prior work has indicated that structural brain connectivity constrains pairwise correlations in brain dynamics (also called functional connectivity), but it is not known whether inter-regional axonal connectivity is related to the intrinsic dynamics of individual brain areas. Here we investigate this relationship using a weighted, directed mesoscale mouse connectome from the Allen Mouse Brain Connectivity Atlas and resting state functional MRI (rs-fMRI) time-series data measured in 184 brain regions in eighteen anesthetized mice. For each brain region, we measured degree, betweenness, and clustering coefficient from weighted and unweighted, and directed and undirected versions of the connectome. We then characterized the univariate rs-fMRI dynamics at each brain region by computing 6 930 time-series properties using the time-series analysis toolbox, *hctsa*. After correcting for regional volume variations, strong and robust correlations between structural connectivity properties and rs-fMRI dynamics were found only when edge weights were accounted for, and were associated with variations in the autocorrelation properties of the rs-fMRI signal. The strongest relationships were found for weighted in-degree, which was positively correlated to the autocorrelation of fMRI time series at time lag *τ* = 34s (partial Spearman correlation *ρ* = 0.58), as well as a range of related measures such as relative high frequency power (*f* > 0.4 Hz: *ρ*= −0.43). Our results indicate that the topology of inter-regional axonal connections of the mouse brain is closely related to intrinsic, spontaneous dynamics such that regions with a greater aggregate strength of incoming projections display longer timescales of activity fluctuations.

---

Nervous systems are complex networks with a topology governed by the pattern of axonal connections linking distinct neural elements. Highly connected and topologically central elements are thought to play an important role in meditating the flow of information across different parts of the system. However, it is unclear how the intrinsic dynamics of a given neuronal population relates to the pattern of connections that population shares with other network nodes. In this work, we show that there is a strong and robust correlation between the structural connectivity properties of a brain region and its blood-oxygenation-level-dependent (BOLD) signal dynamics, as measured with resting-state fMRI (rs-fMRI) in the mouse. The strongest relationship is found with the total weight of incoming connections to a brain region, or weighted in-degree, which is associated with longer dynamical timescales of rs-fMRI dynamics. Our ndings indicate that structural connection weights convey important information about neural activity, and that the aggregate strength of incoming projections to a brain region is closely related to its BOLD signal dynamics.

## I. INTRODUCTION

The principle that structure constrains function is ubiquitous in biology. For example, the molecular structure of a protein determines the species with which it can interact. Similarly, the evolution of opposable thumbs in some primate species enabled high-precision motor control. Brains are no exception, with neuronal dynamics unfolding on an intricate and topologically complex network of axonal connections; a network that is commonly referred to as a connectome^1^. In a graph representation of this network, nodes comprise functionally homogeneous or anatomically localized neurons or populations of neurons (depending on the scale of measurement), and axonal connections between these neural elements are represented as edges connecting pairs of nodes.

The network representation of the brain has provided a convenient framework for understanding the relationship between connectome structure and brain dynamics. This relationship has typically been examined at the level of inter-regional structural connectivity and inter-regional coupling of brain dynamics, or functional connectivity. Correlations between structural and functional connectivity have been demonstrated using a range of approaches and datasets^2–8^, with computational modeling playing a key role. Computational models of brain networks typically simulate dynamical systems (which define the dynamics of each brain region) coupled via a network topology determined by the structural connectome^4–6,9–12^. Some models can predict empirical measurements of functional connectivity in human with model predictions correlating with empirical data in the range 0.4 < *r* < 0.6^13^, and can be optimized up to *r* = 0.75^14^. These results are impressive given the known limitations of diffusion MRI in reconstructing anatomical brain connections^7,15^. The success of dynamical systems models, as well as simplified network spreading models^15–17^, in reproducing the correlation structure of inter-regional brain dynamics suggests that the structural connectome plays a key role in constraining brain dynamics.

While there is a growing evidence base linking the structural topology of a brain network to the inter-regional coupling of functional connectivity, less is known about how connectome structure relates to the intrinsic dynamics of an individual brain region. Understanding this relationship would provide insight into how patterns of neuronal activity within a specific brain area may support its specialized function. In addition to inter-regional variation in microstructural properties and gene transcription^18,19^, it has long been thought that the functional specialization of a given brain region is in large part determined by its unique profile of axonal inputs and outputs - its so-called connectional fingerprint^20^. Moreover, recent work using magnetoencephalogaphy (MEG) has suggested that the dynamics of individual brain regions (captured using power spectral estimates through time) are sufficiently distinctive to be predicted across individuals^21^. Other evidence indicates that brain dynamics are governed by a hierarchy of intrinsic timescales across regions, from slowly-varying prefrontal areas high in the anatomical hierarchy^22^ (that are thought to accumulate information over longer durations), to the relatively rapid dynamics of sensory regions low in the hierarchy^23–28^. This hierarchical organization of timescales across the brain may facilitate processing of (and predictions about) the diverse timescales of stimuli in the world around us. Computational modeling has begun to shed light on the role of connectivity in shaping this inter-regional heterogeneity in characteristic timescales^11^, including the emergence of slow oscillations in densely connected, high-degree brain network hubs in identical, connectome-coupled neural mass models^29^. Thus, although preliminary modeling work has suggested that the connectome may play a role in shaping patterns of dynamical heterogeneity across the brain, empirical data has been lacking to allow a characterization of the relationship between structural connectivity and dynamics at the level of individual brain regions.

Compared to measures of pairwise correlation between time series that yield estimates of functional connectivity, a key challenge of analyzing the univariate dynamics of individual brain regions is the vast array of properties that can be estimated for a given time series recording of neuronal activity. Previous analysis of univariate fMRI dynamics has focused on properties of the power spectrum, such as the total power in particular frequency bands^30^. However, quite apart from properties of the power spectrum, thousands of alternative time series analysis methods exist that might contain useful information, such as those developed for applications in statistics, electrical engineering, economics, statistical physics, dynamical systems, and biomedicine^31^. Here we leverage this vast interdisciplinary library of time series analysis methods to characterize the fluctuations of spontaneous regional activity using resting-state fMRI (rs-fMRI), using a recently developed highly comparative analysis framework that extracts over 7 700 properties from univariate time series^31,32^. In this way, we computed thousands of properties of the intrinsic rs-fMRI dynamics in each brain region using data from 18 anesthetized mice. Structural network connectivity properties of each brain region were also computed using a mesoscale mouse connectome estimated from viral tract-tracing experiments^33^. Comparing the two measurements while correcting for confounding variations in region volumes, we demonstrate robust correlations between a brain region’s structural connectivity and dynamics, with the strongest relationship found between weighted in-degree and autocorrelation properties of the signal. Our results are consistent with the idea that the structural connectome constrains regional rs-fMRI dynamics and underline the importance of measuring weighted connectomes for probing the structure-function relationship.

## II. DATA AND METHODS

Our approach for relating structural connectivity to regional rs-fMRI dynamics is shown schematically in Fig. 1. It involves: (i) extracting topological measures from each node in the network from the structural connectome; (ii) extracting dynamical properties from the fMRI time series measured in each brain region; and (iii) correlating each network property to each dynamical feature. In this section, we first summarize the structural connectivity and functional MRI data used in this study, and then detail the methods used for each of the above steps.

**FIG. 1.**
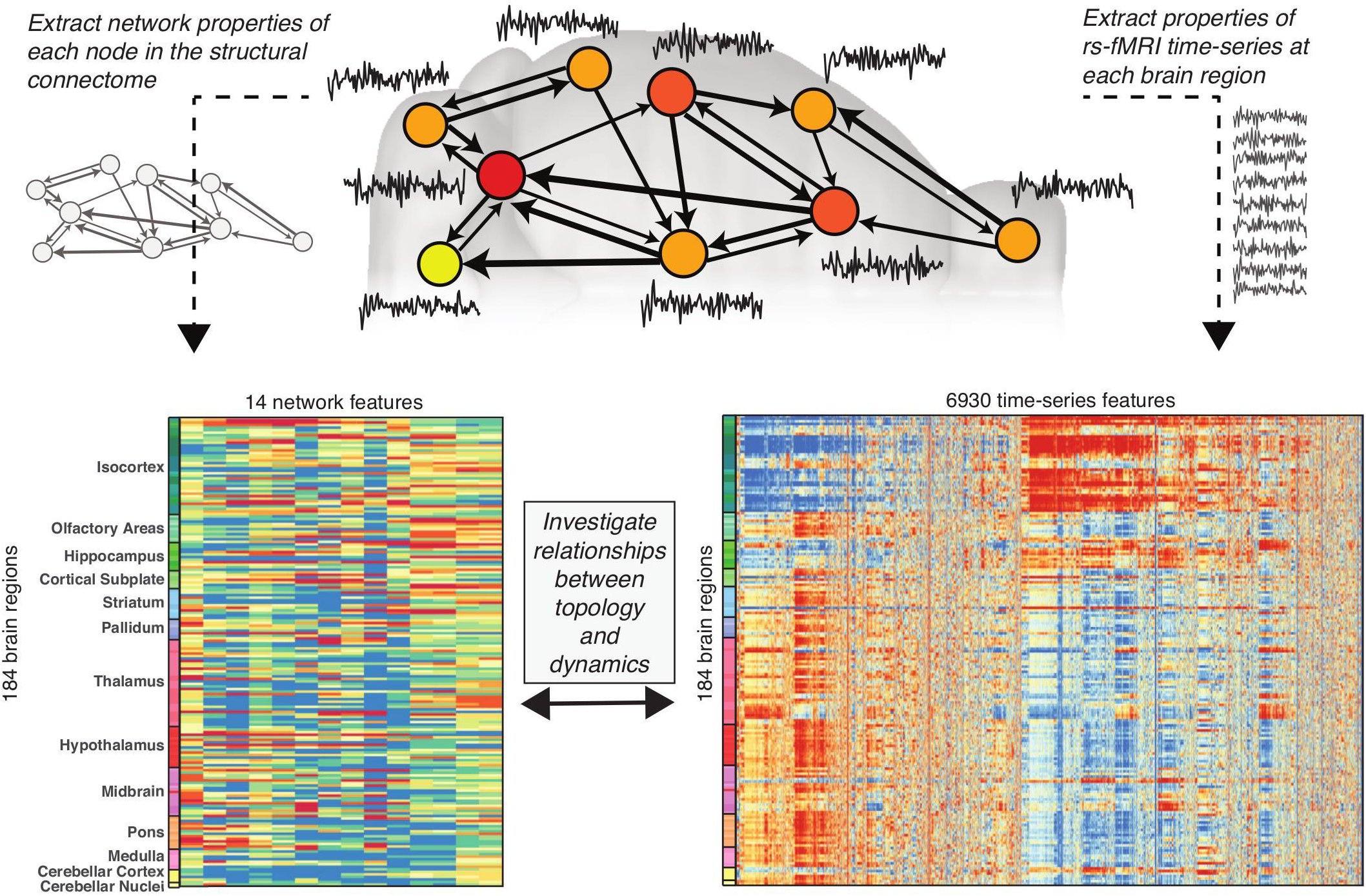
Relating inter-regional connection topological to intrinsic regional dynamics. A schematic illustration of the mouse structural connectome (top), in which the brain is represented as a set of nodes (macroscopic brain regions), with weighted axonal connections between regions represented as directed edges (shown as arrows). different properties can then be computed for each brain region according to their network connectivity properties; the example plotted here is number of connections projecting out of a brain region, also known as ‘out degree’ (shown using color from low, yellow, to high, red). Resting state BOLD dynamics were measured for each brain region using fMRI (shown as time series). Here we compute 14 different network properties for 184 brain regions from the mesoscale structural connectome (lower, left) and, independently, compute 6 930 time-series properties of the univariate fMRI dynamics measured in the same set of brain regions (lower, right). In these lower plots, each row represents a brain region (labeled by broad anatomical divisions of the mouse brain^33,34^), and each column represents a property computed for all brain regions, derived from either the structural connectome (lower, left), or the BOLD time-series dynamics (lower, right). Color encodes the output of each property, from low values (blue) to high values (red). The aim of this study was to determine whether the fMRI dynamics of a brain region are related to its structural connectivity properties by computing correlations between the two quantities across the brain.

### A. Functional MRI data

#### a. Mice

All experiments were performed in accordance to the Swiss federal guidelines for the use of animals in research, and under a license from the Zürich Cantonal veterinary office. Animals were caged in stan-brain. dard housing, with food and water ad libitum, and 12 h day and night cycle.

#### b. Magnetic resonance imaging

Eighteen C57BL/6J mice (age P57 ±7) were used for this experiment. During the MRI session, the levels of anesthesia and mouse physiological parameters were monitored following an established protocol to obtain a reliable measurement of functional connectivity^35^. Briefly, anesthesia was induced with 4% isoflurane and the animals were endotracheally intubated and the tail vein cannulated. Mice were positioned on a MRI-compatible cradle, and artificially ventilated at 80 breaths per minute, 1:4 O_2_ to air ratio, and 1.8 ml/h flow (CWE, Ardmore, USA). A bolus injection of medetomidine 0.05 mg/kg and pancuronium bromide 0.2 mg/kg was administered, and isoflurane was reduced to 1.5%. After 5 min, an infusion of medetomidine 0.1 mg/kg/h and pancuronium bromide 0.4 mg/kg/h was administered, and isourane was further reduced to 0.5%. The animal temperature was monitored using a rectal thermometer probe, and maintained at 36.5 ± 0.5°C during the measurements. The preparation of the animals did not exceed 20 minutes. After the scans, mice were kept under observation in a temperature-controlled chamber with mechanical ventilation (isourane-only, 1%) in order to fully recover from the muscle relaxant agent, the effects of which may last longer than the anesthetic. All animals fully recovered after 30 minutes from the end of the experiment and transferred back to their own cages. Data acquisition was performed on a Pharmascan 7.0 small animal MR system operating at 300 MHz (Bruker BioSpin MRI, Ettlingen, Germany). A high SNR receive-only cryogenic coil (Bruker BioSpin AG, Fallanden, Switzerland) is used in combination with a linearly polarized room temperature volume resonator for transmission. Images were acquired using Paravision 6 software. After standard adjustments, shim gradients were optimized using mapshim protocol, with an ellipsoid reference volume covering the whole brain. Resting-state fMRI (rs-fMRI) was performed with gradient-echo echo planar images (GE-EPI) that were acquired with repetition time TR = 1000ms, echo time TE = 15ms, flip angle = 60°, matrix size = 90 × 50, in-plane resolution = 0.22 × 0.2mm^2^, number of slice = 20, slice thickness ST = 0.4mm, slice gap SG = 0.1mm, 2000 volumes, for a total scan time of 38 min. Anatomical T1-weighted images were acquired with same orientation as the GE-EPI using a FLASH-T1 sequence (TE = 3.51ms, TR = 522ms, flip angle = 30°, in-plane resolution = 0.05 × 0.02 mm^2^, ST = 0.5 mm).

#### c. Data preprocessing

Resting state fMRI datasets were preprocessed using an existing pipeline for removal of unwanted confounds from the time-series^35^, with modifications (Fig. S1). Brifley, each rs-fMRI dataset was fed into MELODIC (Multivariate Exploratory Linear Optimized Decomposition of Independent Components^36^) to perform within-subject spatial-ICA with a fixed dimensionality estimation (number of components set to 100). This included high-pass filtering (> 0.01 Hz), correction for head motion using MCFLIRT^37^ and in-plane smoothing with a 0.3 × 0.3 mm kernel. We applied FSL-FIX with a study-specific classifier obtained from an independent dataset of 15 mice and used a ‘conservative’ removal of the variance of the artifactual components (for more details, see^38^). Thereafter, FIX-cleaned datasets were co-registered into the skull-stripped T1-weighted images and normalized into AMBMC template (http://www.imaging.org.au/AMBMC) using ANTs v2:1 (picsl.upenn.edu/ANTS). Time series were extracted from 370 anatomical regions using the Allen Reference Atlas ontology^34^, as in Oh et al. ^33^. Only regions that were fully covered by the field of view used for fMRI acquisition were included in the analysis. These regions were then matched to the Allen Mouse Connectivity Atlas, above, yielding a total of 184 matching brain regions for each hemisphere. Here we focus on regions in the right hemisphere, for which full structural connectivity data is available (see above). Thus, the final rs-fMRI dataset consisted of 2 000-sample time series in 184 brain regions for 18 mice; a total of 3 312 time series.

### B. Structural connectivity data

The mesoscale structural connectome of the mouse brain was derived from 469 viral microinjection experiments in the right hemisphere of C57BL/6J male mice at age P56, obtained from the Allen Mouse Brain Connectivity Atlas (AMBCA)^33^. These data were summarized in the form of a weighted, directed connectivity matrix containing 213 brain regions from the Allen Reference Atlas ontology^34^ using a regression model^33^. The model outputs a normalized connection strength and a *p*-value for each edge in the connectome, which can be used to construct a 213 × 213 ipsilateral connectivity matrix. We include only edges with *p* < 0.05 (and exclude self-connections), resulting in a link density of 6.9%. Note that our results are not sensitive to this choice; similar results were obtained from denser connectomes derived using higher *p*-value thresholds (up to *p* = 1, an edge density of 37.4%).

The ‘normalized connection strength’ edge weight, estimated directly from the regression model of Oh et al. ^33^, scales the injection volume in a source region to explain the segmented projection volume in a target region. Alternative edge weight measures can be used, which rescale these weights to normalize for the volume of source and target regions^33^, including: ‘connection strength’ (multiplies each edge by the source region’s volume, yielding edge values proportional to the number of axonal fibers projecting from the source to target regions), ‘normalized connection density’ (multiplies each edge by the target region’s volume, yielding edges that measure the fraction of infected volume in the target region resulting from infection of a unit voxel of the source region; used in Rubinov et al. ^39^), and ‘connection density’ (multiplies each edge by the source region’s volume and divides by the target region’s volume, yielding edges that measure the fraction of target region’s volume that would be infected from the entire source region), as visualized in Fig. S2. Given the different interpretation of each edge measure, we compared results using each edge weight definition.

We focus here on results using the full ipsilateral connectome, but also tested the robustness of the results when including contralateral connectivity from the right hemisphere → left hemisphere^33^. From contralateral connectivity data, we inferred a complete connectome under the assumption of hemispheric symmetry (as Rubinov et al. ^39^), in which connections from the left to the right hemisphere match those from right to left hemisphere exactly, and ipsilateral connectivity within the left hemisphere mirrors that within the right hemisphere.

### C. Topological node measures

To characterize the connectivity of each brain region, we used the 213-node, ipsilateral connectome described above to calculate node properties (note that, due to data availability, only 184 of these brain regions were subsequently matched to rs-fMRI dynamics). To assess the role of edge weights and edge directionality, we compared four different representations of networks, where possible: (i) the original directed, weighted connectome(see above); (ii) a directed, unweighted connectome; (iii) an undirected, weighted connectome; and (iv) an undirected, unweighted connectome. For weighted measures, we compared edge weights estimated using each of the different normalizations of the source and target region volumes: the ‘normalized connection strength’, ‘connection density’, ‘connection strength’, and ‘normalized connection density’ (Fig. S2). To transform weighted networks to unweighted networks, we assigned a unit weight to all edges with non-zero weight; to transform directed networks to undirected networks, we assigned edge weights as the sum of the edge weights in and out of each node in the original network. We computed three topological properties for each node in each network: (i) degree; (ii) betweenness centrality; and (iii) clustering coefficient. These measures are described in turn below.

#### d. Degree

For a directed, unweighted network, the in-degree, *k*_in_(*i*), of node *i* is defined as the number of incoming edges, and the out-degree, *k*_out_(*i*), is defined as the number of outgoing edges. On undirected networks, the lack of directional information means that only the total degree, *k*(*i*), can be computed, which is defined as the total number of connections involving node *i*. On weighted networks, the concept of degree can be extended to incorporate edge weights, where the weighted counterpart of node degree (also known as ‘node strength’) sums the weights on edges rather than counting them.

#### e. Betweenness centrality

The betweenness centrality of a node, *i*, is given by

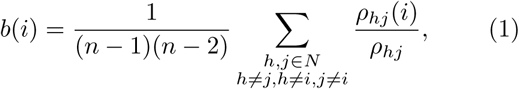

where *N* is the set of all nodes in the network, *n* is the number of nodes, *ρ_hj_* is the total number of shortest paths between nodes *h* and *j*, and *ρ_hj_* (*i*) is the number of those paths that pass through node *i*
^40^. For shortest path information transfer, a node with high betweenness centrality is involved in mediating more signal traffic across the network. For a binary network, all edges have the same weight, and the shortest path between nodes *h* and *j* is the path that minimizes the number of edges that must be traversed. In a weighted network, a distance metric is defined for each link as the inverse of the edge weight.

#### f. Clustering coefficient

The clustering coefficient of a node, *i*, in an unweighted undirected network is given by

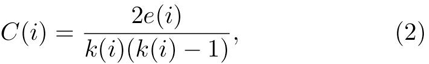

where *k*(*i*) is the degree of node *i* and *e*(*i*) is the number of connected pairs between all neighbors of node *i*^41^. The clustering coefficient of node *i* is equivalent to the link density of its neighbors, such that *C*(*i*) = 1 indicates that node *i* and its neighbors form a clique, i.e., a fully connected subgraph. On an unweighted, directed network, the clustering coefficient of node *i* is defined similarly as *C*^→^(*i*) = *e*(*i*)/[*k*(*i*)(*k*(*i* − 1)]. Weighted generalizations of the clustering coefficient aim to capture the average intensity with which the neighbors of a node are connected. For weighted undirected networks, we use the measure given by Onnela et al. ^42^, and for weighted directed networks we use the measure in Fagiolo ^43^.

For a given edge weight definition, we computed a total of fourteen topological measures for each node: in-degree, out-degree, betweenness and clustering coefficient on the directed weighted and unweighted networks, and degree, betweenness and clustering coefficient on the undirected weighted and unweighted networks. All measures were calculated using implementations provided in the Brain Connectivity Toolbox^44^ (functions used are listed in Supplementary Table S1).

### D. Feature-based representation of rs-fMRI time series

Having quantified different network connectivity properties, we next aimed to characterize the rs-fMRI dynamics in each brain region. BOLD time series have commonly been summarized using *features* like the amplitude of low-frequency (0.01-0.08 Hz) fluctuation, ALFF^30^. Although spectral properties like ALFF are a natural representation of stationary oscillatory signals (as is often approximately the case in brain recordings), there are thousands of alternative time-series analysis methods that could be used to meaningfully quantify regional rs-fMRI dynamics. These methods include measures of autocorrelation, temporal entropy, distributional spread, outlier properties, stationarity, wavelet transforms, time-delay embeddings, and fits to various time-series models. Rather than manually selecting a small number of such time-series features, here we aimed to determine the most informative features for understanding structural connectivity properties in a purely data-driven way. To achieve this, we used the highly comparative time-series analysis software package, *hctsa* (v0.91, github.com/benfulcher/hctsa)^31,32,45^ to extract a to-tal of 7 754 informative features from each of the 3 312 BOLD time series in our dataset (cf. Fig. 1, lower right). Each of the 7 754 features corresponds to a single interpretable measure of a regional BOLD time series, that could then be related to structural network connectivity properties.

Features that did not return a real number for all 3 312 time series in the full dataset (e.g., methods that relied on fitting positive-only distributions to our real-valued rs-fMRI data or methods attempting to fit complicated nonlinear time-series models that were not appropriate for these data), or features that returned an approximately constant value across the dataset (standard deviation < 2 × 10^−15^) were removed from the set of features prior to analysis, resulting in a reduced set of 6 930 well-behaved time-series features. The results of the massive feature extraction facilitated by *hctsa*, was repre-sented as a 184 (brain region) × 6930 (time-series feature) matrix that summarizes a diverse array of BOLD time-series properties in each mesoscale brain region for each mouse. To obtain a group-level summary, we averaged features across all 18 mice for each brain region, yielding an 184 × 6 930 (brain region) × (time-series feature) matrix, which is plotted in Fig. 1 (lower, right). This ensured that the features capture overall characteristics of the BOLD signal at each brain region, averaging over inter-individual and inter-scan variability. In addition to obtaining group-averaged results, we also investigated robustness at the level of each individual mouse by separately analyzing time-series feature matrices for each individual mouse (i.e., 18 different 184 × 6930 matrices).

### E. Relating regional connectivity to rs-fMRI dynamics

Having characterized each brain region in terms of its (i) structural connectivity properties, and (ii) BOLD time series, we next sought to find correlations between these two independent measurements. Spearman rank correlations were used (instead of Pearson correlations) to allow statistical comparison between the frequently non-normally distributed nodal properties (especially those derived from weighted connectomes) and time-series features. To control the family-wise error rate, we used the Holm-Bonferroni method^46^, correcting across 6 930 independent tests at a significance level of *p*_corrected_ < 0.05

Structural connectivity properties and time-series features are both strongly affected by the potentially confounding effect of region volume. We estimated region volume by counting the number of 1.2 *μ*m^3^ isotropic voxels assigned to each brain region; volumes vary markedly across 184 regions, from 17 voxels (subparafascicular area, SPA, in the thalamus) to 5 052 voxels (caudop-utamen, CP, in the striatum). A total of 3 688 time-series features of the rs-fMRI signal were significantly correlated with region volume (*p*_corrected_ < 0.05), with the strongest correlations obtained for spread-dependent measures for which larger regions exhibit reduced variance (e.g., standard deviation: *ρ* = −0.71). Region volume was also related to measures of time-series entropy, spectral power, stationarity, and information theory-based properties. Topological measures were also correlated with region volume for all edge weight definitions; e.g., for ‘connection strength’, weighted topological measures all exhibited correlations *ρ* > 0.5, reaching up to *ρ* = 0.87 for weighted in-degree, as shown in Fig. S3. To ensure that our results reflect the contribution of structural connectivity, and are not a consequence of regional variations volume, we computed partial Spearman rank correlation coefficients, *ρ* (and an associated *p*-value) when comparing connectivity properties to time-series features.

The high quality of these data allowed us to estimate the relationship between nodal network connectivity and rs-fMRI time-series properties using mass univariate testing with family-wise error correction. We correct for 6 930 independent tests, even though there are only approximately 200 linearly independent time-series behaviors in our feature set due to the existence of sets of highly correlated time-series features, cf. Fulcher et al. ^31^. Our results thus constitute a highly conservative estimate of the number of time-series features that are significantly related to each topological quantity, minimizing the false positive rate (type I error) at the cost of artificially inat-ing the false negative rate (type II error). In the absence of such a strong signal, future work could use multivariate methods (such as canonical correlation analysis or partial least squares) to find informative component-wise relationships between the two types of data. This may be especially useful when using connectomes estimated using MRI, which lack directed information and are noisier than the tract tracing-based connectomes analyzed here.

## III. RESULTS

We present our results in three parts. First, we compare nodal network properties derived from weighted/unweighted and directed/undirected connectomes, in their (partial) correlations to the properties of rs-fMRI dynamics across the brain (correcting for region volume). We show that robust correlations between structural network topology and dynamics exist at the level of individual brain regions for weighted connectivity measures, with the strongest relationship found for weighted in-degree, 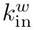. We go on to characterize the correlation properties of rs-fMRI time-series that are most strongly correlated to 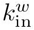. Lastly, we demonstrate that the group-level correlations also hold for the majority of individual mice.

### A. Comparison of topological measures

We first address the question of whether the extrinsic axonal connectivity of a brain region is related to its intrinsic rs-fMRI dynamics. We computed fourteen structural network measures: degree, betweenness, and clustering coefficient, computed from weighted and un-weighted, and directed and undirected versions of the connectome, from each of 184 brain regions. We independently computed 6 930 time-series features derived from the rs-fMRI BOLD signal in the same set of brain regions. The relationship between each structural network property and each rs-fMRI time-series property was quantified as a partial Spearman correlation coefficient (using region volume as a covariate), as depicted in Fig. 1 (see Methods for details).

The comparisons using weighted connectome properties were repeated for each of the four connectome edge weight definitions (depicted in Fig. S2): connection strength, connection density, normalized connection strength, and normalized connection density, as shown in Fig. S4. Although all definitions exhibit qualitatively similar trends, the strongest correlations between rs-fMRI dynamics and topological structure were found when connection weights were proportional to the number of axons connecting two regions (i.e., using ‘connection strength’ or ‘connection density’), i.e., providing a meaningful measure of axonal bandwidth. In the remainder of this study, we focus on results using connection strength edge weights.

Results summarizing the relationship between each structural network property and rs-fMRI dynamics are shown in Fig. 2(a). For each network measure, the figure shows: (i) the magnitude of the strongest partial Spearman correlation, |*ρ*|, across all 6 930 rs-fMRI time-series features (color); and (ii) the number of time-series features that exhibit a significant correlation, (Holm-Bonferroni *p*_corrected_ < 0.05; text annotations). Note that since in- and out-degree cannot be computed from undirected networks; rectangular boxes in the upper right hand quarter of Fig. 2(a) indicate results for degree: *k* (unweighted) and *k*^*w*^ (weighted).

**FIG. 2.**
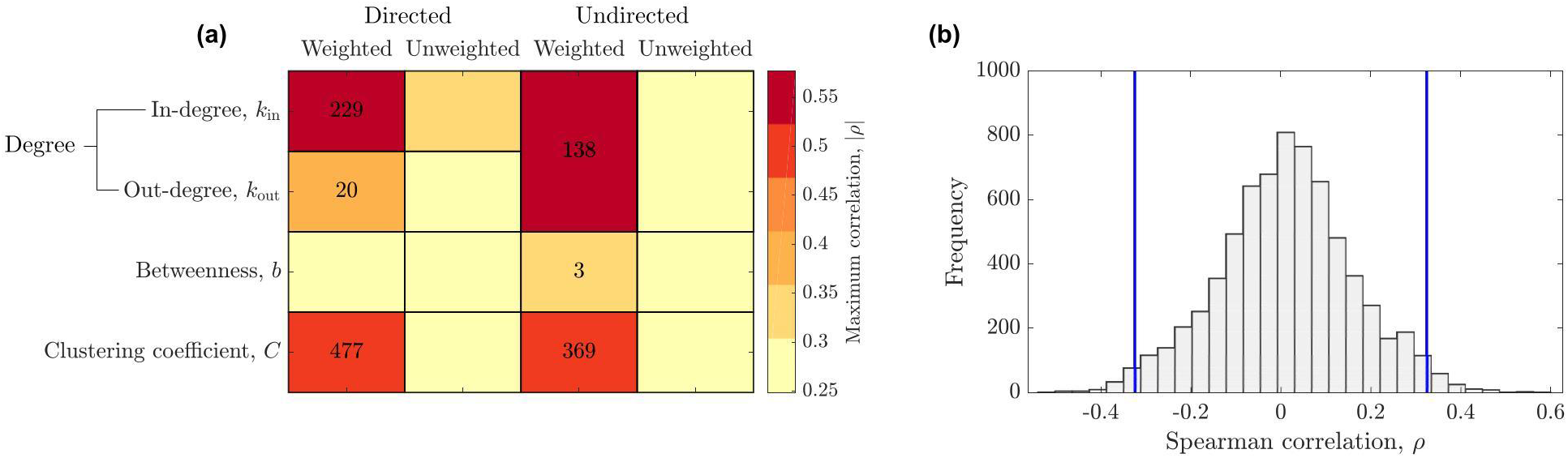
Regional structural network connectivity properties correlate with properties of regional rs-fMRI dynamics. (**a**) We compare the degree, *k*, betweenness, *b*, and clustering coefficient, *C*, for (i) directed and undirected, and (ii) weighted and unweighted structural brain networks (where weights represent connection strengths between regions). For each nodal network property, we computed the magnitude of the strongest partial Spearman correlation, |*ρ*|, controlling for region volume, across 6 930 time-series features of the rs-fMRI signal (shown using color), and the number of time-series features that are signifiantly correlated with that property (Holm-Bonferonni *p*_corrected_ < 0.05) across all 184 brain regions (annotated with numbers; missing numbers indicate zero significant features). We see that taking edge weights into account is crucial for obtaining a strong correlation between regional connectivity and dynamics. (**b**) The distribution of Spearman correlations, *ρ* of weighted in-degree, 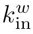, with 6 930 time-series features of rs-fMRI (correlations computed across 184 brain regions). Vertical blue lines indicate Holm-Bonferonni significance thresholds at *p*_corrected_ = 0.05.

Although we summarize the results using the maximum correlation, max(|*ρ*|), in Fig. 2(a), the comparison of each network property to 6 930 rs-fMRI time series properties is best represented as a distribution of correlations, such as that shown in Fig. 2(b) for weighted in-degree, 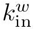. Figure 2(b) indicates the thresholds for Holm-Bonferroni *p*_corrected_ < 0.05 (vertical blue lines), revealing a large number of rs-fMRI properties that correlate strongly and significantly with 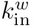 across 184 mouse brain regions, with correlations reaching up to |*ρ*| = 0.58. Similar distributions for all node measures are in Fig. S5.

Connection strengths vary over many orders of magnitude, from a connection strength of 0:15 for the weakest connection to 7.03 ×10^3^ for the strongest connection (arbitrary units, cf. Fig. S6). Node-level structural network properties are therefore highly sensitive to the incorporation of edge weights. Strong and robust correlations to time-series properties were only found for network properties derived from weighted connectomes, whereas unweighted connectome measures exhibited weak correlations after controlling for region volume (|*ρ*| < 0:31), with none exhibiting statistically significant correlations after Holm-Bonferroni correction. Amongst the weighted measures, most connectivity properties exhibit stronger correlations to rs-fMRI dynamics when edge directionality was taken into account, pointing to an important role of edge directionality for uncovering the relationship between structural connectivity and dynamics.

Of the three nodal structural connectome properties analyzed here, the immediate measure of connectivity, degree, *k*, showed the strongest correlations to regional rs-fMRI dynamics. While a significant correlation was found using the weighted total degree, *k*^*w*^ (up to |*ρ*| = 0.53), when distinguishing incoming and outgoing connection pathways, our results reveal an asymmetry, with weighted in-degree, 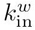 showing the strongest correlations to rs-fMRI dynamics of all topological properties (up to |*ρ*| = 0.58), while weighted out-degree, 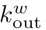, showed weaker correlations (up to |*ρ*| = 0.39). This increase in correlation for 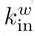 over 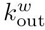 demonstrates the relative importance of incoming structural connectivity for understanding regional BOLD dynamics (and this trend holds across all edge weight definitions, cf. Fig. S4). In addition to degree, we found significant but weaker correlations between clustering coefficient and properties of the rs-fMRI dynamics. In the directed network, *C*^→, *w*^ significantly correlated with 477 time-series features (*p*_corrected_ < 0.05, with correlations reaching as high as |*ρ*| = 0.51), while *C*^*w*^ computed from the undirected network was significantly correlated with 369 time-series features (up to |*ρ*| = 0.48). Note that the number of significant features depends on the shape of the distribution (like that shown in Fig. 2), and that the maximal partial correlation, |*ρ*|, of any single topological measure does not necessarily reflect the total number of features with *p*_corrected_ < 0.05. Betweenness centrality was the least correlated nodal connectivity property to properties of rs-fMRI dynamics (a result that was consistent across all edge weight definitions, cf. Fig. S4). Only three of the 6 930 time-series features were significantly correlated with weighted undirected betweenness, *b*^*w*^, (up to |*ρ*|= 0.33) and there were no significant correlations for *b*^→, *w*^, *b*^*w*^, or *b*.

Noting that network connectivity measures are themselves non-independent and interrelated (perhaps due to redundancy in the space of allowed networks^?^), we calculated Spearman rank correlation coefficients between all pairs of the fourteen topological measures, across all 184 brain regions (Fig. S7). Topological measures that showed the strongest partial correlations with time-series features 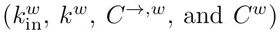 formed a strongly in-tercorrelated group (|*ρ*| > 0.7 for all pairs), while the unweighted measures *C*^→^, *C*, *b*^→^, *b* and *k*, were also strongly intercorrelated. The high level of intercorrelation between 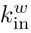, *k*^*w*^, *C*^→,*w*^, and *C*^*w*^ suggests that they are measuring similar connectivity properties in different ways, and may therefore be related to similar properties of the rs-fMRI signal.

Given that weighted in-degree, 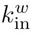 shows the strongest relationship to rs-fMRI dynamical features, and forms a correlated cluster with other types of weighted connectivity properties, we tested the idea that the variation of 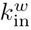 is sufficient to explain the relationship of the other informative weighted connectivity properties: *C*^*w*^, *C*^*→*, *w*^, and *k*^*w*^. For each of these topological measures, we computed partial correlations to all 6 930 rs-fMRI time-series features, controlling for 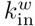 (and region volume). The number of time-series features that were significantly related to these measures dropped dramatically after controlling for 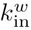 (*p*_corrected_ < 0.05): C^*→*, *w*^ (477 → 4), *C*^*w*^ (369 → 1), and *k*^*w*^ (138 → 0), indicating that the much of the signal relating *C*^*w*^, *C*^→,w^, and *k*^*w*^ with rs-fMRI dynamics can be explained by their correlation with 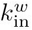 However, it is not the case that 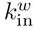 is uniquely capturing information about rs-fMRI time-series. For example, computing a partial correlation between 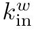 and time-series properties controlling for both region volume and *k*^*w*^ reduces the number of significant features (*p*_corrected_ < 0.05) to just two; controlling for *C*^*→*, *w*^ reduces the number to 25; controlling for all other topological quantities together reduces the number to zero. Thus, although it does not capture distinct information from other connectivity properties, 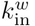 exhibits the strongest correlation to time-series features, with a variation across brain regions that can mostly account for the relationship of other topological quantities to rs-fMRI properties.

### B. Informative time-series features

Having demonstrated a strong relationship between weighted structural connectivity properties of a brain region and its rs-fMRI dynamics, and that weighted in degree, 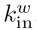, shows the strongest correlation, we next characterize the types of rs-fMRI time-series properties that drive the effect, focusing on this key topological measure, 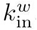. The 229 rs-fMRI time-that were significantly correlated with 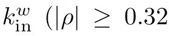, *p*_corrected_ < 0.05) constitute different measures of autocorrelation properties of the fMRI signal. For example, of the five features with partial Spearman correlations |*ρ*| > 0.5 (correcting for region volume variations), four were direct measures of autocorrelation computed at extended timescales (*τ* = 16,23,26,34 s), with the top feature being autocorrelation at *τ* = 34 s: AC_34_ (partial Spearman correlation, *ρ* = 0.58). Other related features include nonlinear autocorrelations (e.g., 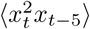, with the average taken across the time series, *x*: *ρ* = 0:46), automutual information (e.g., Gaussian estimator at time lag *τ* = 34 s: *ρ* = 0.44), power in spectral frequency bands (e.g., power in the range of frequencies from 0.125 Hz ≤ *f* ≤ 0.375 Hz: *ρ* = −0.48), parameters of model fits (e.g., a_5_ parameter of an autoregressive AR(5) model: *ρ* = −0.51), measures of randomness (e.g., *p*-value from an Ljung-Box Q-test^?^: *ρ* = −0.39), and entropy (e.g., normalized permutation entropy, PermEn(*m* = 3, *τ* = 2)^?^: *ρ* = −0.36). Between themselves, these features have highly correlated outputs across brain regions, indicating that they are measuring similar properties of the data in different ways (Fig. S8). Thus, using a highly comparative data-driven approach to univariate time-series analysis^31,32,45^, our results indicate that regions with increased 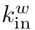 (relative to their volume) exhibit increased autocorrelation, which can be measured directly, or with related measures, such as power in specific frequency bands, or some classes of entropy/predictability measures.

To investigate the variation in these time-series features across brain regions, we first focus on the feature correlating most strongly with 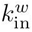, autocorrelation at a lag of *τ* = 34s (AC_34_: *ρ* = 0.58). The rank residuals of both 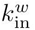, and AC_34_ (after correcting for region volume using a partial Spearman correlation) are plotted as a scatter in Fig. 3(a). The clear positive trend (*ρ* = 0.58), indicates that brain regions with a larger weighted in-degree exhibit stronger autocorrelations at this extended, 34 s timescale.

Given the ubiquity of power spectral analysis in neuroimaging, we next focus on a related spectral feature, which measures relative high-frequency power, *f* ≥ 0.4 Hz (up to the Nyquist frequency, 0.5 Hz), and correlates strongly with 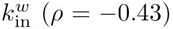. The power spectral density of the BOLD signal is estimated using a periodogram with a Hamming window applied, and this feature calculates the logarithm of the proportion of the power in the top fifth of frequencies (i.e., *f* ≥ 0.4 Hz), which we term ‘relative high frequency power’ (where ‘high’ is the range from 0.4 Hz up to the Nyquist frequency, 0.5 Hz). This feature displays a negative cor-relation with 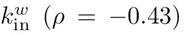 across the brain, shown as a scatter in Fig. 3(b). That is, brain regions with greater 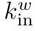 have decreased relative power in rs-fMRI dynamics for *f* ≥ 0.4 Hz (or, equivalently, have greater relative power in *f* < 0.4 Hz). This is consistent with intuition from AC_34_, which increases with 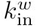; brain regions with increased aggregate incoming connection weights exhibit ‘slower’ BOLD dynamics. To demonstrate the relationship in more detail, we plotted power spectral density estimates for three selected brain regions in Fig. 3(b): the magnocellular part of cicular nucleus (SPFm) in the thalamus 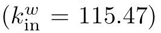, the sensory-related superior colliculus (SCs) in the mid-brain 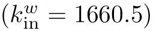, and the ventral part of the anterior cingulate area (ACAv) in the isocortex 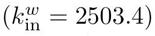, as annotated in Fig. 3(c). Frequencies exceeding *f* = 0.4 Hz are shaded in Fig. 3(c), indicating the decrease in high frequency power in regions with a higher 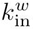.

**FIG. 3.**
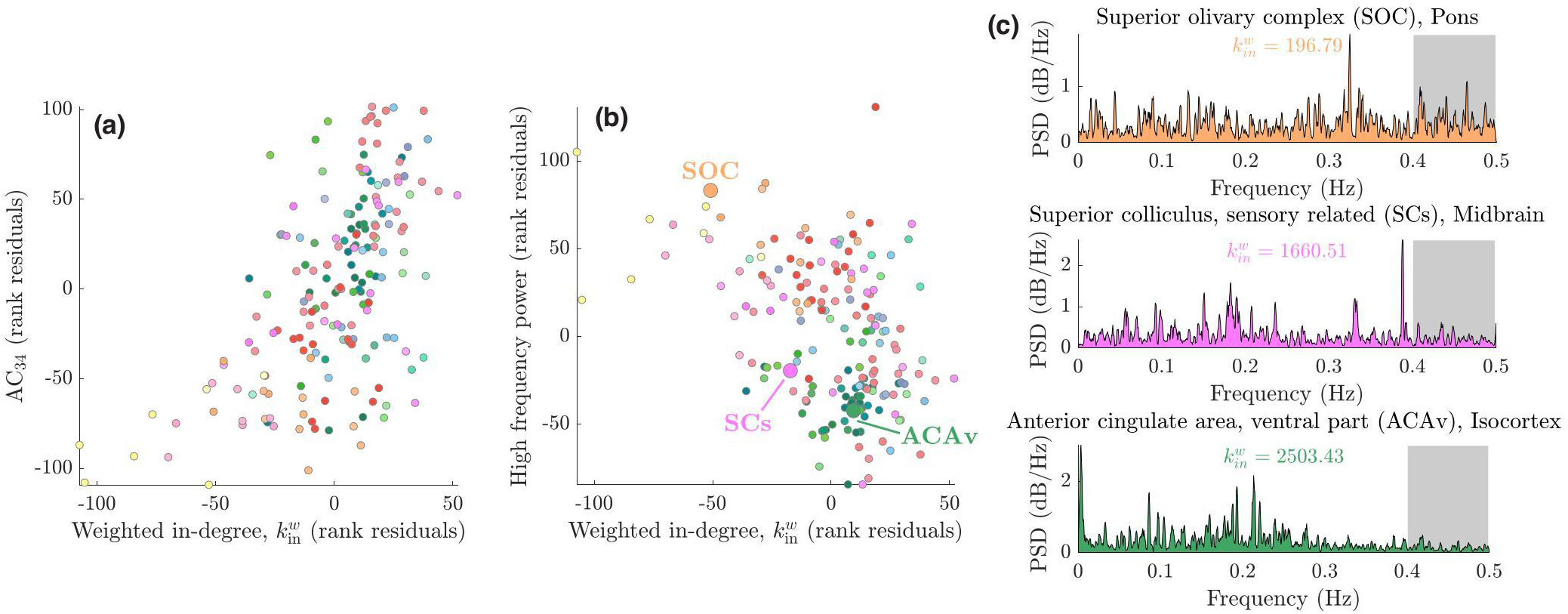
Weighted in-degree is negatively correlated with regional BOLD signal relative high-frequency power. Scatter plots of a brain region’s weighted in-degree are shown against (**a**) autocorrelation of the signal at *τ* = 34s, and (b) relative power of the rs-fMRI signal in frequencies *f* ≥ 0.4 Hz. For each variable rank residuals are plotted, after correcting for a Spearman correlation to region volume. Brain regions are colored uniquely, corresponding to anatomical divisions (cf. Fig. 1). For three selected brain regions, highlighted and labeled in (**b**), we show power spectral density estimates of their rs-fMRI signals calculated using a periodogram (smoothed for visualization) in (**c**). The relative high frequency power measured in (**b**) corresponds to the logarithm of the proportion of the power in the top fifth of frequencies (shaded gray) of the power spectral density estimate, as calculated using a periodogram. This corresponds to the frequency range *f* ≥ 0.4 Hz, and is lower for regions with increased 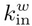.

As noted above, some dynamical properties of the rs-fMRI signal differ across anatomical divisions (as is evident by the visual distinction of particularly the isocortex and hippocampus in Fig. 1, lower right), and may thus result from broad anatomical differences, or non-specific spatial gradients in dynamics, rather than reflecting specific properties of network connectivity. We mapped the spatial variation of weighted in-degree, 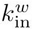, AC_34_, and relative high frequency power (as Spearman rank residuals after correcting for region volume variations) across the brain to better visualize their spatial variation, as shown in Fig. 4. The rendering for 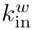 (residuals) in Fig. 4(a), shows a specific distribution across the brain, with peaks in the ventral striatum region (i.e., nucleus accumbens, NAc), a region that integrates a large number of cortical and midbrain neural inputs, in the superior colliculus (SC), a subcortical area that integrates visual and sensory information, in the thalamus and in the cornu ammonis 1 region of the hippocampus (CA1), which is involved in memory and learning. A similar spatial specificity is reflected in the variation of rs-fMRI AC_34_ residuals across the brain in Fig. 4(b). The high-frequency power feature [inverted in Fig. 4(c)] is high in olfactory cortex, midbrain and cerebellar regions, and low in isocortex, caudoputamen and thalamus.

**FIG. 4.**
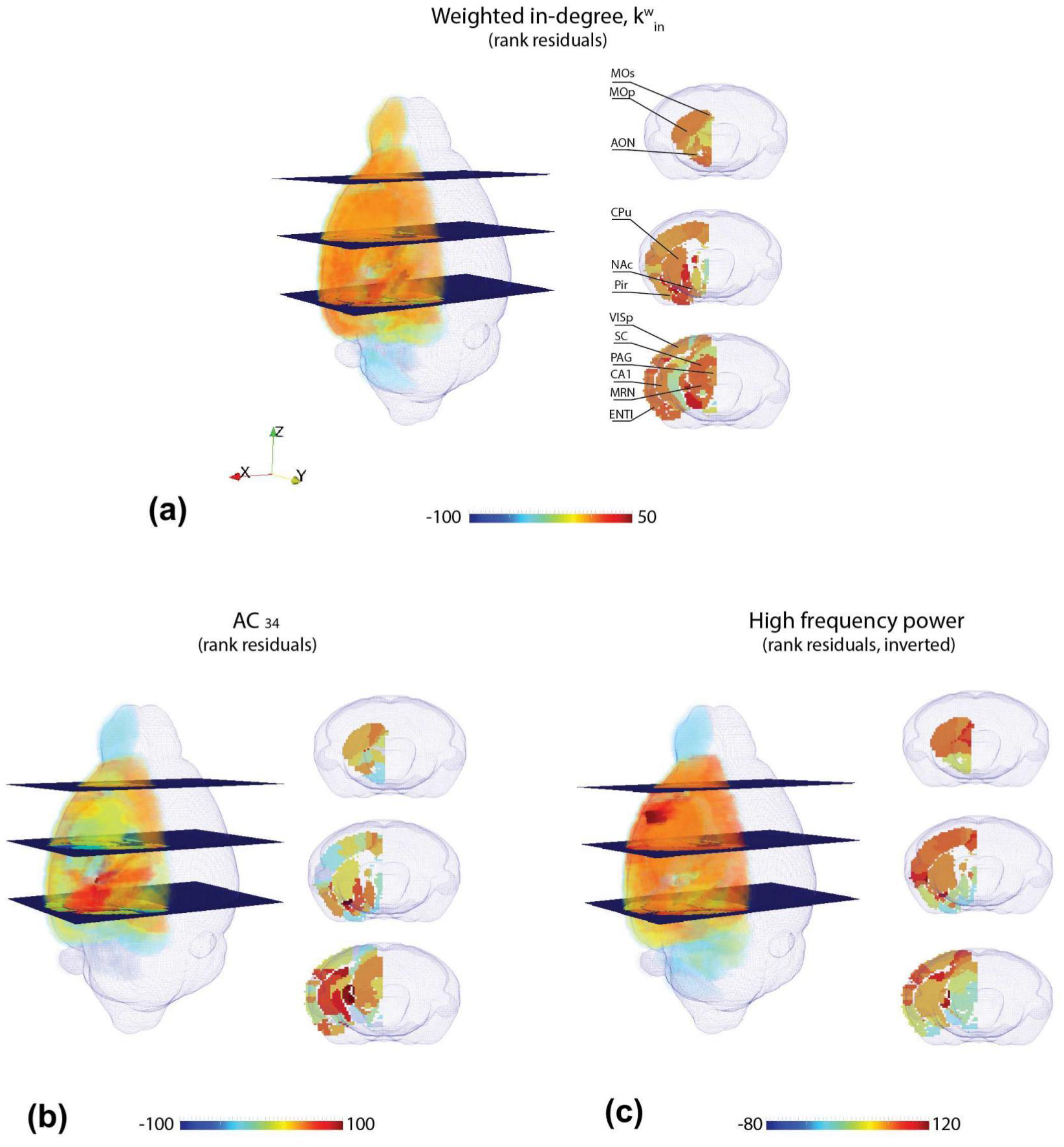
A three-dimensional rendering of the spatial mapping of 184 regions across the right hemisphere of the mouse brain for: (**a**) the topological quantity, weighted in-degree, 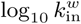, (**b**) autocorrelation at lag *τ* = 34 s, and (**c**) relative high frequency BOLD power (*f* ≥ 0.4 Hz, note inverted color scale). All above variables have been plotted as the rank residuals from a Spearman partial correlation with region volume. Labelled regions in (a) are: MOp, MOs = primary and secondary motor cortex; AON = anterior olfactory nucleus; CPu = Caudoputamen; NAc = nucleus accumbens; Pir = piriform cortex; VISp = primary visual area; SC = superior colliculus; PAG = periaqueductal gray; CA1 = cornus ammonis 1; MRN = midbrain reticular nucleus; ENTI = entorhinal area.

Above, we focused on characterizing the time-series features that strongly correlate with 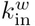 with the expectation that similar types of properties would be selected for the other significant connectivity properties that are highly correlated to 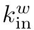, namely, *k*^*w*^, *C*^→,*w*^, and *C*^*w*^ [see Fig. 2(a)]. Most of the rs-fMRI time-series features that were significantly correlated to *k*^*w*^, *C*^→,*w*^ and *C*^*w*^, were subsets of the 229 features selected for 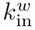, with all selected features related to linear or nonlinear autocorrelations and power spectral properties.

Our analysis above was of ipsilateral connectivity in the complete connectome available in the right hemisphere of the mouse brain. Inclusion of contralateral projections, under the assumption of hemispheric symmetry (see Methods), yielded qualitatively similar results to those presented here. In particular, 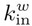 remained the topological property with the strongest relationship to rs-fMRI dynamics, and exhibited similar correlations to autocorrelation-based measures of the rs-fMRI signal, including the AC_34_ (*ρ* = 0:49, *p*_corrected_ = 9 × 10^−9^) and relative high frequency power (*ρ* = − 0.36, *p*_corrected_ = 0.003) features described above.

### C. Individual robustness

The results above involved characterizing the rs-fMRI dynamics of each brain region by averaging time-series features across all 18 individual mice. Here we analyze the relationships at the level of individual mice. The strongest correlating feature, the autocorrelation of the BOLD signal at a time lag *τ* = 34s (AC_34_), showed a group-level correlation with weighted in-degree, 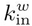, of *ρ* = 0.58. Computing this relationship for each individual mouse yielded partial Spearman correlations ranging between −0.22 ≤ *ρ* ≤ 0.55, with a significant relationship found for 13 of the 18 mice (right-tailed partial Spearman correlations: *p*_corrected_ < 0.05, Holm-Bonferroni corrected for 18 comparisons). A similar analysis was performed for relative high frequency power (*f* ≥ 0.4 Hz, group-level *ρ* = −0.43), with Spearman correlation coefficients ranging from −0.64 ≤ *ρ* ≤ −0.14 across the 18 individual mice. A significant correlation was observed for each of the 18 mice (*p*_corrected_ < 0.05), with 15 of the 18 mice exhibiting a stronger partial correlation than at the group level (i.e., *ρ* < −0.43). Although there is variability across individual mice, these results indicate that the group-level relationships are not a consequence of averaging over a group of mice, but hold for the majority of individual mice.

## IV. DISCUSSION

In this work we used a weighted, directed mesoscale structural connectome and high quality rs-fMRI measurements across 184 anatomical brain regions to demonstrate a robust relationship between a brain region’s topological role in the structural connectome and its resting state dynamics in the mouse. Rather than analyzing pairwise structure-function relationships, our results characterize a potential role of structural connectivity in shaping the dynamics of individual brain regions. We show that weighted connectivity information is required to uncover the regional structure-function relationship, and that the weight of incoming projections to a region is a key correlate for regional BOLD dynamics, particularly with respect to its autocorrelation (and related spectral power and other measures). As well as providing new insights into potential functional implications of structural brain connectivity, our empirical results may provide a new means of constraining the models we use to simulate and understand brain dynamics.

We first note the strong effect of region volume variations on both rs-fMRI dynamics and connectivity measures. The number of inter-areal axonal connections is expected to scale with the volume of brain areas and, consequently, other properties derived from the weighted connectome (cf. Fig. S3). Similarly, the strong relationship found between a large number of time-series properties of rs-fMRI dynamics is partially accounted for by image processing steps that involve averaging over different numbers of voxels to produce an overall BOLD time series for a given brain region. We found that larger brain regions had lower standard deviation and reduced high frequency power. However, although corrected for here using partial Spearman correlations, region volume is unlikely to be a pure confound, as the mean parcellation size varies markedly across the brain, from very small average region sizes in the thalamus, medulla, and pons, for example, and moderate to larger volumes in the isocortex, olfactory areas, and hippocampal formation. By computing partial correlations that correct fully for variation in areal volumes, we may therefore be conservative in the effects we report, as this correction may remove some signal of connectivity-dynamical coupling, due to the properties of the anatomical parcellation used.

After correcting for the volume of brain areas, the strongest correlations between structural connectivity and rs-fMRI dynamics were found for measures of immediate connectivity (degree) and neighborhood connectivity (clustering coeffcient), with the strongest individual correlation found for weighted in-degree, 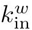. By contrast, the global measure of betweenness centrality showed minimal correlations to dynamics (potentially related to the fact that information transmission across brain networks may more closely follow an unguided, diffusion-like process rather than shortest paths^6,15,16^). Weighted in-degree was significantly correlated to 229 rs-fMRI time-series properties (*p*_corrected_ < 0.05, from a set of 6 930 features), with partial Spearman correlation coefficients reaching up to |*ρ*| = 0.58 (for linear autocorrelation at lag *τ* = 34s). Given the diversity of regions across the whole mouse brain analyzed here, that differ in their functional specialization, gene expression, and cellular and microcircuit properties^18,19^, all of which may affect regional dynamics^11,47^, this level of correlation with just the incoming mesoscale connectivity to a brain region, 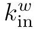, is remarkable. Apart from 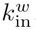, all other network properties that showed strong and significant correlations to rs-fMRI dynamics—*k*^*w*^, *C*^*w*^, and *C*^→,*w*^—were related to similar types of time-series properties as 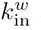 and showed minimal correlations to rs-fMRI dynamics when 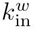 was controlled for. Taken together, our findings indicate that direct influences from other areas, as measured by 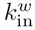 (or highly correlated measures *k*^*w*^, *C*^*w*^, and *C*^→,*w*^) are closely tied to a brain region’s spontaneous dynamics. Our findings may also reflect the hemodynamic measure of neuronal activity provided by the BOLD signal. The strongest neurophysiological correlate of the BOLD signal is the local eld potential, which is more strongly driven by local synaptic integration of incoming signals rather than spiking output^48,49^. Whether the close association between incoming connectivity and dynamics reported here for mesoscopic brain regions would also hold using single unit recordings of individual neurons remains an open question.

Rather than hand-picking particular time-series features to investigate for rs-fMRI, our highly comparative approach compared over 6 930 time-series features of fMRI in a purely data-driven way, including time-series model fitting and prediction, entropies, distributional measures, and other types of linear and nonlinear correlation features^31,32,45^. Of the 229 properties of regional rs-fMRI dynamics that were significantly corre-lated to weighted in-degree, 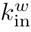, those with the strongest relationship were measures of autocorrelation, including power in spectral frequency bands, parameters of linear models, and entropy measures. Regions with increased 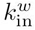 were most strongly characterized by the autocorrelation of their BOLD dynamics at a time lag *τ* = 34s (*ρ* = 0.58). A range of autocorrelations at similar lags also exhibited strong correlations to weighted in-degree; e.g., the four features with the highest correlation to 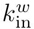 were all autocorrelations at different time lags: AC_34_ (*ρ* = 0.58), AC_26_ (*ρ* = 0.56), AC_23_ (*ρ* = 0.53), and AC_16_ (*ρ* = 0.52). To be selected in a purely data-driven way from 6 930 features, this result suggests an importance of these relatively long timescales (lags between 16s and 34s). Other features measured similar autocorrelation properties of the signal in less direct ways, such as computing spectral power in the upper fifth of sampled frequencies, *f* ≥ 0.4 Hz (*ρ* = −0.43). These group-level effects were robust, being reproduced at the level of 13 of the 18 mice for AC_34_ (*p*_corrected_ < 0.05), and 17 of the 18 individual mice for relative high frequency power (*p*_corrected_ < 0.05).

Our results are consistent with a connectivity-mediated hierarchy of timescales in the brain that has been suggested by some computational models^11,29^. In Gollo et al. ^29^, nonlinear neural mass models were coupled via an unweighted, directed macaque structural connectome, producing an emergent dynamical hierarchy in which highly connected hub regions exhibited greater temporal persistence in their dynamics, largely due to their high (unweighted) in-degree. In Chaudhuri et al. ^11^, an interplay of intrinsic variation in timescales across cortical brain regions, inter-areal connectivity, and input to the brain determined the dynamical timescale of a brain region (estimated as the decay time constant of the autocorrelation function). Although these models provide candidate mechanisms that may explain variations in fMRI autocorrelation timescales across regions of the mouse brain, it remains unclear whether this relationship is due to the presence of incoming connections directly altering the dynamics of that brain region. While this is a compelling possibility, our results could also be explained by a common underlying factor, with other types of well-studied heterogeneity in cytoarchitecture and gene expression giving rise both to differences in inter-regional connectivity profiles and characteristic BOLD dynamics, for example. Future work will be crucial to disentangling the causal mechanisms underlying the correlational relationships presented here.

Our tract-tracing derived mouse connectivity data allowed us to investigate the role of edge weights and directionality in contributing to the relationship between the structural connectome and fMRI dynamics. In particular, we compared measures computed from the weighted and directed connectome^33^ (comparing different edge weight definitions), as well as unweighted and undirected approximations of it. Although unweighted brain networks are more intuitive and amenable to the application of graph theoretic techniques that have traditionally been popular in network neuroscience, one might expect that incorporating meaningful estimates of edge weights into brain network analysis is important, as they vary over several orders of magnitude (see Fig. S6). Indeed, given recent estimates of cortical connection densities from high-quality tract-tracing data exceeding 60% for macaque cortex^50,51^ and 70% for mouse cortex^52^, binary representations of such dense connectomes constitute coarse approximations of true brain connectivity. Here we demonstrate that network properties derived from unweighted connectomes do not show strong correlations to univariate rs-fMRI properties, highlighting the importance of measuring connectome edge weights in capturing the relationship between connectivity and dynamics in the mouse brain. We note that the estimation of edge weights from tract-tracing based experiments is non-trivial; here we explored different edge weight definitions derived from the regression model of Oh et al. ^33^, but note that alternative estimation methods^39,52^ and datasets^53^ exist. Of the four edge weight definitions considered here (that normalize source and target regional volumes differently, cf. Fig. S2), those that took into account the source volume (i.e., ‘connection strength’ and ‘connection density’, which take into account the volume of each source region) yielded weighted connectomes that showed the strongest correlations to rs-fMRI dynamics. Interpreting this with respect to weighted in-degree, 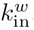, for example, this indicates that the connectome representations that provide the most information about dynamics in a target region are those in which edge weights account for the volume of source regions that project it, i.e., obtain a measure proportional to the number of projecting axons. Indeed, the strongest relationships to rs-fMRI dynamics were found when connectome edge weights were proportional to the number of axonal pathways between two regions (i.e., using ‘connection strength’). Given that human connectomes are commonly estimated by tractography using diffusion weighted imaging (DWI), which is noisy and cannot determine the direction of a pathway, it is important to note that *k*^*w*^ is highly correlated to 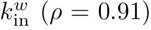, and is related to similar types of features of rs-fMRI dynamics as 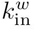 (most features are a subset of those selected for 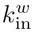, and all are correlated with a feature selected for 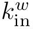 with *ρ* > 0.59). This suggests that, if similar connectivity-dynamics relationships hold in human as in mouse, then weighted degree, *k*^*w*^, calculated from an undirected connectome should also contain meaningful information about autocorrelation properties of rs-fMRI dynamics. Although weighted degree, *k*^*w*^, measured from the undirected connectome showed strong correlations to rs-fMRI dynamics (up to |*ρ*| = 0.53), directed edge information was important for distinguishing the weighted out-degree of a node, 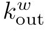, which is relatively uninformative of rs-fMRI dynamics (up to |*ρ*| = 0.39), from the most informative weighted in-degree, 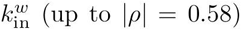 Taking edge weight and directionality into account when relating structure to function in brain networks will be crucial to understanding whether brain dynamics are causally constrained by extrinsic axonal connectivity in future work.

The mouse represents an attractive model system to study the structure-function relationship, while minimizing the influence of environmental and genetic heterogeneity. Our findings relied on long (38 min), high-quality rs-fMRI measurements of BOLD dynamics across the whole mouse brain, which were compared to the a tract-tracing based structural connectome for the first time in this work. The use of awake mice in rs-fMRI protocols is impracticable for long scan times (notwithstanding the use of invasive methods for head fixation^54^), making light anesthesia the de facto option^55^. As previously demonstrated in rats^56^ and monkeys^57^, decreasing (or abolishing) levels of anesthesia are mirrored by an increased in BOLD variability. This indicates that data acquired during anesthesia cannot be fully generalized to the awake status. Yet, to minimize the effects of anesthesia on BOLD dynamics, we employed a combination of low-dose isoflurane+medetomidine, which previously demonstrated its efficiency in retaining strong bilateral brain networks connectivity (i.e., one of the signature of rs-fMRI observed in both awake humans and rodents)^58^.

Due to the technical and methodological challenges in obtaining such long functional scans in a light-anesthetized regime, one must take care that the physiology of the animals remains stable over time. Several parameters were considered in this study to achieve this goal; first, we used mechanical ventilation to maintain the same tidal volume and blood oxygenation throughout the experiment. To keep a low but steady level of anesthesia, we combined a continue infusion of medetomedine i/v with a very low dose of isoflurane (0.5%), optimized from our previous studies^35,59^; this allowed us to overcome the issues related to medetomidine depletion and isoflurane accumulation over time. As evidenced by our results, which showed consistency across the majority of individual mice, this protocol allows for superior data reproducibility due to a drastic reduction in motion, stable physiology such as animal temperature and stress levels, and a regular breathing cycle.

## V. CONCLUSION

In this work, we provide the first demonstration of a robust relationship between the connectivity properties of a brain region and resting state BOLD dynamics. The strongest relationships were observed for the weighted indegree of a brain region, which is positively correlated to autocorrelations at extended time lags (e.g., *τ* = 34s) of its rs-fMRI dynamics and, similarly, negatively correlated to their relative high frequency power (*f* ≥ 0.4 Hz). Our results are consistent with physiological data indicating that BOLD signal modulations reflect the integration of incoming signals, and support preliminary predictions of computational models suggesting a role for connectivity in mediating differences in the intrinsic dynamical timescales of distinct regions across the brain, with highly connected brain regions exhibiting slower BOLD dynamics. These findings also highlight an asymmetry between incoming and outgoing connectivity, and underline the importance of weighted information for understanding brain communication across structural brain networks.

## SUPPLEMENTARY MATERIAL

See supplementary material for additional figures and tables referred to in the text.

## ACKNOWLEDGMENTS

We would like to thank L. Gollo for providing valuable advice and feedback on the manuscript, and the two reviewers for their crucial insights, particularly for prompting us to investigate the influence of inter-regional volume variations. We would like also to acknowledge G. Ielacqua and M. Markicevic for their precious help during the MRI experiments. V.Z. is supported by the ETH Zurich Postdoctoral Fellowship Program and by the ETH Career Seed Grant SEED-42 16-1. A.F. is supported by an ARC Future Fellowship (FT130100589) and NHMRC Project Grants (1066779 and 1050504). B.D.F. is supported by a NHMRC Early Career Fellowship (1089718). The authors declare no competing nancial interests.

## References

1 Sporns O. The human connectome: a complex network. Ann. N.Y. Acad. Sci. 1224, 109 (2011).

2 Deco G., Robinson P. A., Jirsa V. K., Breakspear M. J., and Friston K. J. The Dynamic Brain: From Spiking Neurons to Neural Masses and Cortical Fields. PLoS Comp. Biol. 4, e1000092 (2008).

3 Skudlarski P., Jagannathan K., and Calhoun V. D. Measuring brain connectivity: diffusion tensor imaging validates resting state temporal correlations. NeuroImage (2008).

4 Honey C. J., Sporns O., Cammoun L., et al. Predicting human resting-state functional connectivity from structural connectivity. Proc. Natl. Acad. Sci. USA 106, 2035 (2009).

5 Honey C. J., Thivierge J.-P., and Sporns O. Can structure predict function in the human brain? NeuroImage 52, 766 (2010).

6 Goñi J., van den Heuvel M. P., Avena-Koenigsberger A., et al. Resting-brain functional connectivity predicted by analytic measures of network communication. Proc. Natl. Acad. Sci. USA 111, 833 (2014).

7 Zalesky A. and Fornito A. A DTI-Derived Measure of Cortico-Cortical Connectivity. IEEE Trans. Med. Imaging 28, 1023 (2009).

8 Finger H., Bönstrup M., Cheng B., et al. Modeling of Large-Scale Functional Brain Networks Based on Structural Connectivity from DTI: Comparison with EEG Derived Phase Coupling Networks and Evaluation of Alternative Methods along the Modeling Path. PLoS Comp. Biol. 12, e1005025 (2016).

9 Honey C. J., Breakspear M., Kötter R., and Sporns O. Network structure of cerebral cortex shapes functional connectivity on multiple time scales. Proc. Natl. Acad. Sci. USA 104, 10240 (2007).

10 Hagmann P., Deco G., Ponce-Alvarez A., et al. Resting-State Functional Connectivity Emerges from Structurally and Dynamically Shaped Slow Linear Fluctuations. J. Neurosci. 33, 11239 (2013).

11 Chaudhuri R., Knoblauch K., Gariel M.-A., Kennedy H., and Wang X.-J. A Large-Scale Circuit Mechanism for Hierarchical Dynamical Processing in the Primate Cortex. Neuron 88, 419 (2015).

12 Mišić B., Betzel R. F., de Reus M. A., et al. Network-Level Structure-Function Relationships in Human Neocortex. Cereb. Cortex 26, 3285 (2016).

13 Deco G. and Jirsa V. K. Ongoing Cortical Activity at Rest: Criticality, Multistability, and Ghost Attractors. J. Neurosci. 32, 3366 (2012).

14 Deco G., McIntosh A. R., Shen K., et al. Identi cation of optimal structural connectivity using functional connectivity and neural modeling. J. Neurosci. 34, 7910 (2014).

15 Mišić B., Betzel R. F., Nematzadeh A., et al. Cooperative and Competitive Spreading Dynamics on the Human Connectome. Neuron 86, 1518 (2015).

16 Goñi J., Avena-Koenigsberger A., de Mendizabal N. V., et al. Exploring the Morphospace of Communication E ciency in Complex Networks. PLoS ONE 8 (2013).

17 Mišić B., Sporns O., and McIntosh A. R. Communication Ef-ciency and Congestion of Signal Traffic in Large-Scale Brain Networks. PLoS Comp. Biol. 10, e1003427 (2014).

18 Lein E., Hawrylycz M. J., Ao N., et al. Genome-wide atlas of gene expression in the adult mouse brain. Nature 445, 168 (2007).

19 Madisen L., Zwingman T. A., Sunkin S. M., et al. A robust and high-throughput Cre reporting and characterization system for the whole mouse brain. Nat. Neurosci. 13, 133 (2010).

20 Passingham R. E., Stephan K. E., and Kotter R. The anatomical basis of functional localization in the cortex. Nat. Rev. Neurosci. 3, 606 (2002).

21 Keitel A. and Gross J. Individual Human Brain Areas Can Be Identified from Their Characteristic Spectral Activation Fingerprints. PLoS Biol. 14, e1002498 (2016).

22 Felleman D. J. and Van Essen D. C. Distributed Hierarchical Processing in the Primate Cerebral Cortex. Cereb. Cortex 1, 1 (1991).

23 Kiebel S. J., Daunizeau J., and Friston K. J. A Hierarchy of Time-Scales and the Brain. PLoS Comp. Biol. 4, e1000209 (2008).

24 Honey C. J., Thesen T., Donner T. H., et al. Slow Cortical Dynamics and the Accumulation of Information over Long Timescales. Neuron 76, 423 (2012).

25 Stephens G. J., Honey C. J., and Hasson U. A place for time: the spatiotemporal structure of neural dynamics during natural audition. J. Neurophysiol. 110, 2019 (2013).

26 Murray J. D., Bernacchia A., Freedman D. J., et al. A hierarchy of intrinsic timescales across primate cortex. Nat. Neurosci. 17, 1661 (2014).

27 Chen J., Hasson U., and Honey C. J. Processing Timescales as an Organizing Principle for Primate Cortex. Neuron 88, 244 (2015).

28 Cocchi L., Sale M. V., Gollo L. L., et al. A hierarchy of timescales explains distinct effects of local inhibition of primary visual cortex and frontal eye fields. eLife Sciences 5, e15252 (2016).

29 Gollo L. L., Zalesky A., Hutchison R. M., van den Heuvel M. P., and Breakspear M. J. Dwelling quietly in the rich club: brain network determinants of slow cortical fluctuations. Phil. Trans. Roy. Soc. B 370, 20140165 (2015).

30 Yu-Feng Z., Yong H., Chao-Zhe Z., et al. Altered baseline brain activity in children with ADHD revealed by resting-state functional MRI. Brain Dev. 29, 83 (2007).

31 Fulcher B. D., Little M. A., and Jones N. S. Highly comparative time-series analysis: the empirical structure of time series and their methods. J. Roy. Soc. Interface 10, 20130048 (2013).

32 Fulcher B. D. and Jones N. S. Automatic time-series phenotyping using massive feature extraction. bioRxiv p. 081463 (2016).

33 Oh S. W., Harris J. A., Ng L., et al. A mesoscale connectome of the mouse brain. Nature 508, 207 (2014).

34 Dong H.-W. The Allen reference atlas: A digital color brain atlas of the C57Bl/6J male mouse. John Wiley & Sons Inc., Hoboken, NJ, US (2008).

35 Zerbi V., Grandjean J., Rudin M., and Wenderoth N. Mapping the mouse brain with rs-fMRI: An optimized pipeline for functional network identi cation. NeuroImage 123, 11 (2015).

36 Beckmann C. F. and Smith S. M. Probabilistic Independent Component Analysis for Functional Magnetic Resonance Imaging. IEEE Trans. Med. Imaging 23, 137 (2004).

37 Jenkinson M., Bannister P., Brady M., and Smith S. Improved Optimization for the Robust and Accurate Linear Registration and Motion Correction of Brain Images. NeuroImage 17, 825 (2002).

38 Gri anti L., Salimi-Khorshidi G., Beckmann C., et al. ICA-based artefact removal and accelerated fMRI acquisition for improved resting state network imaging. NeuroImage 95, 232 (2014).

39 Rubinov M., Ypma R. J. F., Watson C., and Bullmore E. Wiring cost and topological participation of the mouse brain connectome. Proc. Natl. Acad. Sci. USA 112, 10032 (2015).

40 Freeman L. C. A Set of Measures of Centrality Based on Betweenness (1977).

41 Watts D. J. and Strogatz S. H. Collective dynamics of ‘small-world’ networks. Nature 393, 440 (1998).

42 Onnela J.-P., Saramaki J., Kertesz J., and Kaski K. Intensity and coherence of motifs in weighted complex networks. Phys. Rev. E 71, 065103 (2005).

43 Fagiolo G. Clustering in complex directed networks. Phys. Rev. 76 (2007).

44 Rubinov M. and Sporns O. Complex network measures of brain connectivity: Uses and interpretations. NeuroImage 52, 1059 (2010).

45 Fulcher B. D. and Jones N. S. Highly comparative feature-based time-series classi cation. IEEE Trans. Knowl. Data Eng. 26, 3026 (2014).

46 Holm S. A Simple Sequentially Rejective Multiple Test Procedure. Scand. J. Stat. 6, 65 (1979).

47 Marblestone A. H., Wayne G., and Kording K. P. Toward an integration of deep learning and neuroscience. Front. Comput. Neurosci. 10, 406 (2016).

48 Logothetis N. K., Pauls J., Augath M., Trinath T., and Oelter-mann A. Neurophysiological investigation of the basis of the Fmri signal. Nature 412, 150 (2001).

49 Lauritzen M. and Gold L. Brain function and neurophysiologi-cal correlates of signals used in functional neuroimaging. J. Neurosci. 23, 3972 (2003).

50 Markov N. T., Ercsey-Ravasz M. M., and Gomes A. A weighted and directed interareal connectivity matrix for macaque cerebral cortex. Cereb. Cortex (2012).

51 Markov N. T., Ercsey-Ravasz M., Van Essen D. C., et al. Cortical High-Density Counterstream Architectures. Science 342, 1238406 (2013).

52 Ypma R. J. F. and Bullmore E. Statistical Analysis of Tract-Tracing Experiments Demonstrates a Dense, Complex Cortical Network in the Mouse. PLoS Comp. Biol. 12, e1005104 (2016).

53 Zingg B., Hintiryan H., Gou L., et al. Neural Networks of the Mouse Neocortex. Cell 156, 1096 (2014).

54 Yoshida K., Mimura Y., Ishihara R., et al. Physiological effects of a habituation procedure for functional MRI in awake mice using a cryogenic radiofrequency probe. J. Neurosci. Methods 274, 38 (2016).

55 Jonckers E., Delgado y Palacios R., Shah D., et al. different anesthesia regimes modulate the functional connectivity outcome in mice. Magn. Reson. Med. 72, 1103 (2014).

56 Bettinardi R. G., Tort-Colet N., Ruiz-Mejias M., Sanchez-Vives M. V., and Deco G. Gradual emergence of spontaneous correlated brain activity during fading of general anesthesia in rats: Evidences from fmri and local eld potentials. NeuroImage 114, 185 (2015).

57 Barttfeld P., Uhrig L., Sitt J. D., et al. Signature of consciousness in the dynamics of resting-state brain activity. Proceedings of the National Academy of Sciences 112, 887 (2015).

58 Grandjean J., Schroeter A., Batata I., and Rudin M. Optimization of anesthesia protocol for resting-state fmri in mice based on differential effects of anesthetics on functional connectivity patterns. NeuroImage 102, Part 2, 838 (2014).

59 Grandjean J., Schroeter A., Batata I., and Rudin M. Optimization of anesthesia protocol for resting-state fMRI in mice based on differential effects of anesthetics on functional connectivity patterns. Neuroimage 102 Pt 2, 838 (2014).

